# Neuroimaging Supports the Representational Nature of the Earliest Human Engravings

**DOI:** 10.1101/464784

**Authors:** E Mellet, M Salagnon, A Majki, S Cremona, M Joliot, G Jobard, B Mazoyer, Mazoyer N Tzourio, F d’Errico

## Abstract

The earliest human graphic productions dating to the Lower and Middle Palaeolithic are associated with anatomically modern and archaic hominins. These productions, which consist of abstract patterns engraved on a variety of media, may have been used as symbols, and their emergence is thought to be associated with the evolution of the properties of the visual cortex. To test this hypothesis, we used functional magnetic resonance imaging to compare brain activations triggered by the perception of engraved patterns dating between 540,000 and 30,000 years before the present with those elicited by the perception of scenes, objects, symbol-like characters, and written words. The perception of the engravings bilaterally activated regions in the ventral route in a pattern similar to that produced by the perception of objects, suggesting that these graphic productions are processed as organized visual representations in the brain. Moreover, the perception of the engravings led to a leftward activation of the visual word form area. These results support the hypothesis that in contrast to random doodles, the earliest abstract graphic productions had a representational purpose for modern and archaic hominins.

---

From Palaeolithic rock paintings to contemporary art, abstract and figurative graphic productions have represented a major aspect of human cognitive activity. The ability to produce visual representations was assumed to have emerged at the beginning of the Upper Palaeolithic era approximately 42,000 years BP (Before Present) due to a sudden genetic mutation occurring only in anatomically modern human populations1. The subsequent discoveries of abstract paintings and engravings older than 42,000 years in Africa and Eurasia suggest that this ability emerged much earlier and was shared by different fossil species, including *Homo erectus*^2^, Neanderthals^3,4^ and Early *African Homo sapiens*. Analyses of several early abstract patterns have shown that these patterns were produced deliberately and had no apparent utilitarian function^2,3,5,6^, leading some authors^7^ to argue that these ancient abstract patterns represent the earliest instances of symbol use a thousand years prior to the earliest known 40- to 30,000-year-old examples of depictional representations^8^. The production of abstract patterns has been linked to the evolution of the primary visual areas and surrounding cortex^9,10^. Alternatively, the complexification of graphic productions from simple geometric to depictional representations may reflect the successive evolutionary stages of visual information processing through the primary visual cortex to the occipito-temporal cortex via the ventral route. The ventral route is involved in the extraction of visual information and allows the recognition of visual categories, such as shapes, scenes, and words (see^11^ and^12^ for a review). If the abstract patterns intentionally engraved by hominins were not random doodles and were perceived as structured forms with a potential meaning, their perception should engage the ventral route cortex, which extends anteriorly to the early visual areas in the occipito-temporal pathway. We test this hypothesis in the present study, which aims to characterize the cerebral regions involved in the perception of the earliest engraved abstract graphisms (EEAG). The blood oxygen level dependent (BOLD) signal was mapped in twenty-seven healthy volunteers using functional magnetic resonance imaging (fMRI), while the individuals were presented with tracings of EEAG dating between 540 ka and 30 ka (See Table S1). The perception of these patterns was compared to the perception of their scrambled version, in which the genuine organization of the abstract patterns was lost to assess whether the geometric organization of the EEAG (albeit some patterns were very simple) could be differentiated at the visual cortex level from patterns with no perceptual organization. Another aim of the study was to shed light on the nature of the EEAG. Therefore, we assessed whether the regions activated by the perception of EEAG were also involved in the processing of other visual categories visually presented to the same participants, including a scrambled version of each stimulus belonging to each categoriy (Figure 1).

**Figure 1.**
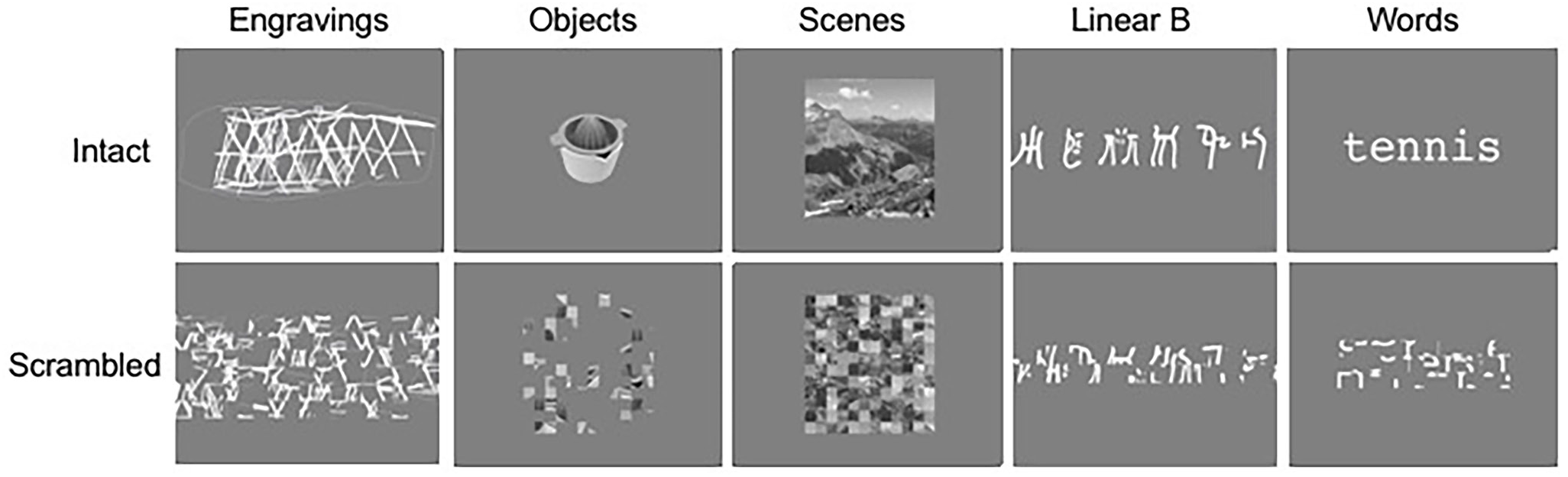
Example of intact and scrambled stimuli used in the 1-back task.

The first category included pictures of nameable human-made artefacts. These stimuli conveyed semantic and lexical information but no symbolic content. The rationale for this choice is that proximity between the patterns of activation of the EEAG and objects could support the representational character of the EEAG, which could also indicate that the EEAG were processed as a whole rather than a collection of lines, fulfilling the “good shape” criteria of the gestalt theory of perception^13,14^. The second category was represented by outdoor scenes depicting largescale surroundings. These stimuli provided information regarding the outer world and conveyed semantic content without being perceived as a single item. In addition, two symbolic categories were included in the study. The first symbolic category included chains of characters from the linear B syllabic script, which is a writing system unknown to the participants, and these characters were chosen as stimuli perceptible as ‘potentially’ symbolic. A similar profile of activation between the EEAG and linear B script stimuli could suggest that the EEAG are processed as symbols or signs that result from a combination of structurally simple elements. The second symbolic category was represented by two-syllable words. This condition extended the requirements of processing the linear B alphabet because the perception of familiar individual overlearned signs (letters) and familiar combinations trigger access to the pronunciation and meaning of words. In addition, word reading is characterized by a leftward asymmetric activation of the ventral route that reflects symbol processing and lexical and semantic access^15,16^. Thus, this condition provides a pattern of leftward asymmetry that could be compared with the asymmetry elicited by the perception of EEAG. The regional BOLD values in response to the engravings *minus* the scramble contrast were extracted from each homotopic region (hROI) of the functional homotopic atlas AICHA^17^. The regions included in the analysis fulfilled the following two criteria: in at least one hemisphere, a significant positive value was observed in response to both the engravings minus scramble contrast (p < 0.05 False discovery rate, FDR corrected, one-sample two-sided t-test) and engravings minus fixation (see methods) contrast (p < 0.05 uncorrected, one- sample two-sided t-test). The fulfilment of these conditions ensured that regions with stronger deactivation under the reference condition (Scramble) during the EEAG comparison with the fixation were not selected.

Ten hROIs were significantly activated in either the right or left hemisphere in the engravings minus scramble contrast. All hROIs were located in the visual ventral pathway (Table 1, Table S2 and Figure 2). Thus, the perceptual processing of EEAG compared to their scrambled version did not involve primary visual areas but did involve visual regions beyond the primary visual and peristriate areas. Previous work has led authors to propose that the early visual cortex may play a role in the extraction of the geometric features composing EEAG^9,10^. However, the present work shows that the activity generated by the perception of scrambled and intact EEAG in the early visual cortex does not substantially differ. This similarity suggests that the evolution of the properties of the primary visual cortex was unlikely to play a central role in the emergence of EEAG or that the latter triggered the former.

**Table 1.** Mean BOLD value of the ten hROIs activated in the engraving minus scramble contrast and left minus right asymmetry under the five conditions. * significant (one-sample test, p < 0.05 corrected).

|  | Olat3 | O2-2 | O3-2 | O3-1 | T3-5 | FUS5 | T3-4 | FUS4 | FUS3 | FUS2 |
| --- | --- | --- | --- | --- | --- | --- | --- | --- | --- | --- |
| **Engravings** |  |  |  |  |  |  |  |  |  |  |
| BOLD LH | 0.25 (0.11)* | 0.16 (0.06)* | 0.46 (0.09)* | 0.49 (0.09)* | 0.27 (0.07)* | 0.20 (0.05)* | 0.31 (0.08)* | 0.39 (0.08)* | 0.18 (0.04)* | 0.13 (0.05)* |
| BOLD RH | 0.45 (0.16)* | 0.17 (0.07)* | 0.43 (0.10)* | 0.36 (0.10)* | 0.16 (0.06)* | 0.25 (0.07)* | 0.19 (0.11)* | 0.28 (0.09)* | 0.13 (0.04)* | 0.12 (0.04)* |
| Asym L-R | -0.20 (0.09) | 0.00 (0.03) | 0.03 (0.06) | 0.13 (0.06) | 0.11 (0.04) | -0.05 (0.04) | 0.13 (0.06) | 0.12 (0.04)* | 0.05 (0.02) | 0.00 (0.03) |
| **Objects** |  |  |  |  |  |  |  |  |  |  |
| BOLD LH | 0.37 (0.08)* | 0.11 (0.05)* | 0.52 (0.07)* | 0.55 (0.08)* | 0.26 (0.05)* | 0.34 (0.04)* | 0.37 (0.07)* | 0.43 (0.07)* | 0.19 (0.04)* | 0.19 (0.03)* |
| BOLD RH | 0.48 (0.10)* | 0.15 (0.05)* | 0.41 (0.06)* | 0.16 (0.07)* | 0.12 (0.05)* | 0.25 (0.04)* | -0.06 (0.07) | 0.25 (0.05)* | 0.10 (0.03)* | 0.17 (0.03)* |
| Asym L-R | -0.11 (0.08) | -0.04 (0.03) | 0.11 (0.05)* | 0.40 (0.07)* | 0.13 (0.04)* | 0.08 (0.03)* | 0.43 (0.04)* | 0.19 (0.05)* | 0.09 (0.02)* | 0.02 (0.03) |
| **Words** |  |  |  |  |  |  |  |  |  |  |
| BOLD LH | -0.22 (0.06)* | -0.41 (0.05)* | -0.28 (0.06)* | -0.47 (0.08)* | -0.22 (0.05)* | -0.18 (0.03)* | -0.08 (0.07) | 0.25 (0.06)* | 0.07 (0.04) | 0.07 (0.03)* |
| BOLD RH | -0.43 (0.08)* | -0.36 (0.06)* | -0.45 (0.07)* | -0.66 (0.06)* | -0.23 (0.04)* | -0.27 (0.04)* | -0.58 (0.08)* | -0.16 (0.05)* | -0.07 (0.03)* | 0.04 (0.03) |
| Asym L-R | 0.21 (0.06)* | -0.04 (0.04) | 0.17 (0.04)* | 0.19 (0.08)* | 0.01 (0.06) | 0.10 (0.04)* | 0.50 (0.08)* | 0.41 (0.04)* | 0.14 (0.03)* | 0.11 (0.03)* |
| **Linear B** |  |  |  |  |  |  |  |  |  |  |
| BOLD LH | 0.32 (0.08)* | -0.02 (0.05) | 0.33 (0.07)* | 0.19 (0.09)* | 0.06 (0.06) | 0.14 (0.04)* | 0.14 (0.07) | 0.40 (0.07)* | 0.14 (0.04)* | 0.14 (0.04)* |
| BOLD RH | 0.42 (0.10)* | 0.01 (0.04) | 0.26 (0.06)* | -0.01 (0.08) | 0.01 (0.06) | 0.04 (0.04) | -0.12 (0.07) | 0.23 (0.04)* | 0.11 (0.03)* | 0.10 (0.03)* |
| Asym L-R | -0.10 (0.09) | -0.03 (0.04) | 0.06 (0.04) | 0.20 (0.06)* | 0.05 (0.05) | 0.10 (0.04)* | 0.25 (0.05)* | 0.17 (0.06)* | 0.03 (0.02) | 0.04 (0.03) |
| **Scenes** |  |  |  |  |  |  |  |  |  |  |
| BOLD LH | -0.02 (0.06) | 0.21 (0.04)* | 0.12 (0.04)* | 0.07 (0.05) | 0.06 (0.03) | 0.31 (0.04)* | 0.07 (0.05) | 0.10 (0.05) | 0.06 (0.03) | 0.07 (0.02)* |
| BOLD RH | 0.14 (0.06)* | 0.20 (0.04)* | 0.15 (0.05)* | 0.00 (0.05) | 0.09 (0.03)* | 0.44 (0.05)* | -0.07 (0.06) | 0.11 (0.04)* | 0.10 (0.03)* | 0.12 (0.02)* |
| Asym L-R | -0.16 (0.03)* | 0.01 (0.03) | -0.04 (0.02) | 0.07 (0.03)* | -0.04 (0.02) | -0.13 (0.04)* | 0.14 (0.03)* | -0.01 (0.03) | -0.04 (0.02) | -0.05 (0.02)* |

**Figure 2.**
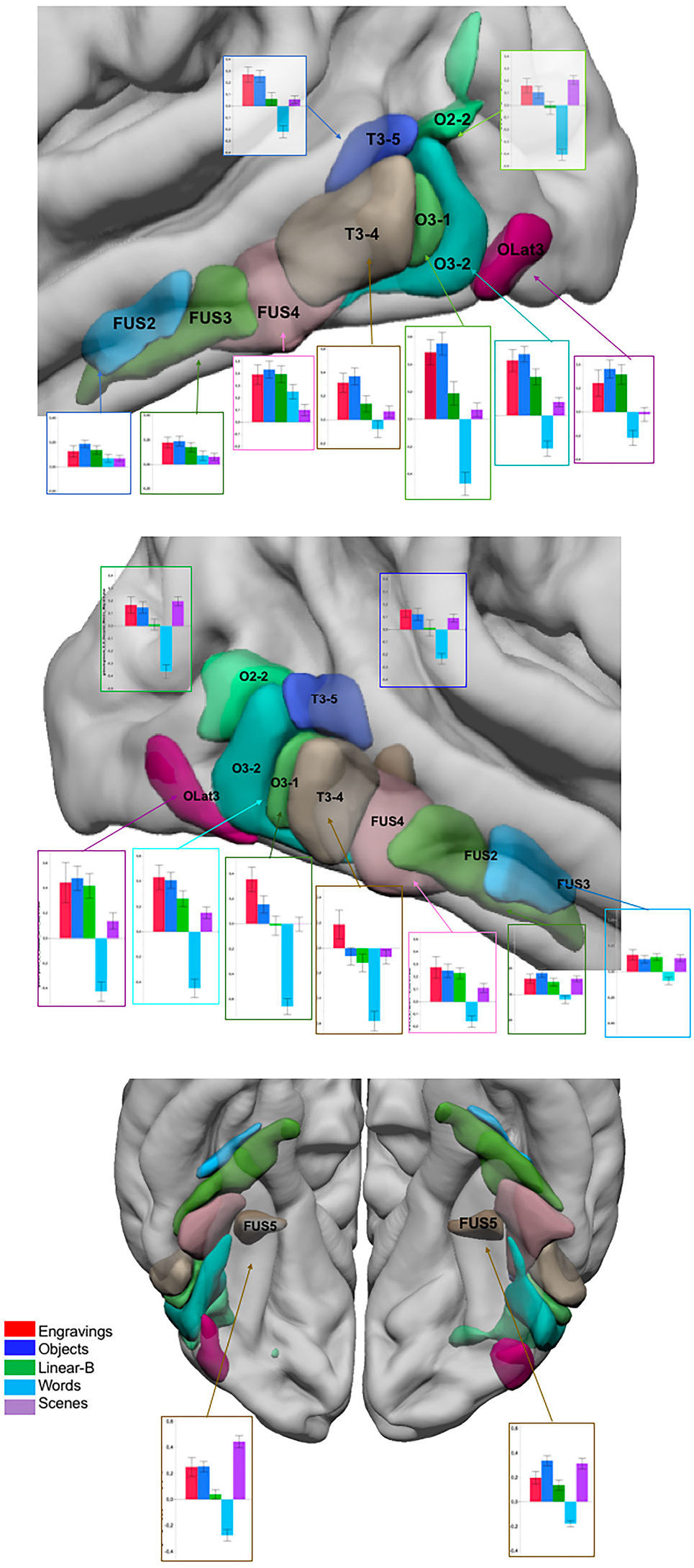
Superimposition on an MRI template of the 10 hROIs showing a significant BOLD signal increase in the engravings *minus* scramble contrast. **Top:** Lateral view of the left hemisphere, **Middle:** Lateral view of the right hemisphere, **Bottom:** Inferior view of the left and right hemispheres. Bar plots represent the BOLD values obtained in the five categories *minus* their scrambled version contrasts in each region. Error bars represent the standard error of the mean.

Although EEAG are not recognized as existing entities, their perception elicited activation in anterior regions of the ventral pathway sensitive to distinct visual categories^11,12,18,19^. Since the EEAG and their scrambled version differed because the latter lacked geometric organization, the level of perceptual organization required for EEAG appears to be sufficiently high to recruit higher order visual areas. The involvement of the visual ventral pathway indicates that information processing requires more than the mere identification of visual primitives, such as the simple extraction of edges, oriented segments, or ends of lines.

Interestingly, in the left hemisphere, the 10 regions activated in the EEAG *minus* scramble contrast were also significantly activated by the perception of objects compared to their scrambled version (Table 1 and Figure 2). Repeated-measures ANOVA comparing EEAG and object perception conditions showed that there was no hROIs x condition interaction effect on the BOLD values (*F*(9,225) = 1.6, p = 0.11), reflecting the similarity of activation between these two conditions in the left hemisphere. In the right hemisphere, a condition by region interaction was present (*F*(9,225) = 2.4, p = 0.02) due to a significantly lower BOLD value in the right inferior temporal hROI (T3-4) under the object condition than under the EEAG condition. No other regions showed a significant BOLD difference between these two conditions in the right hemisphere. The activation profile in response to each visual category compared to its scrambled version among the 10 hROIs confirms that the EEAG and object conditions triggered similar activations along the ventral pathway (Figure 3). Notably, the activation profile under the EEAG perception condition in the 16 hROIs activated by object perception (listed in Table S3 as supplementary material) was also identical (see figure S1 in supplementary material), providing supplementary evidence of the brain functional overlap between object and EEAG perception.

**Figure 3.**
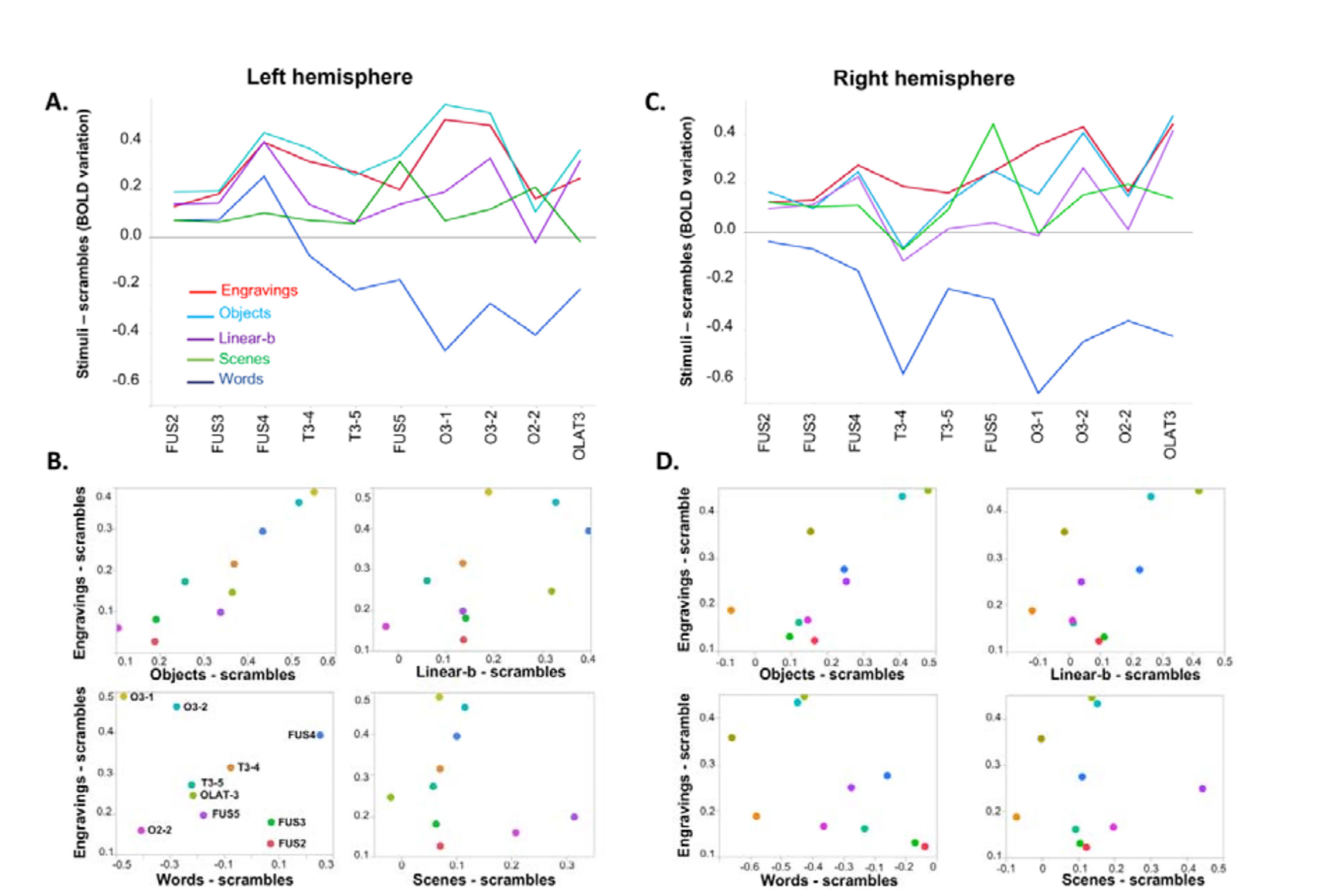
Descriptive analysis of the activation profiles of the 10 hROIs of the ventral route activated by EEAG perception under the different conditions. **Top:** Profiles of activity in the left hemisphere (**A**) and right hemisphere (**C**) in regions presented in an antero-posterior sequence along the ventral route. **Bottom:** Plots of the BOLD values obtained under the EEAG condition against those obtained under the objects, linear B words and scenes conditions across the 10 hROIs in the left (**B**) and right (**D**) hemispheres. Notably, the values on the horizontal axis can be positive, null, or negative according to the contrast and the hROIs.

In contrast, the comparison of the BOLD values observed under the EEAG condition with those in each of the other conditions revealed strongly significant hROIs x conditions interactions in all cases in both the left [EEAG, LinearB: *F*(9,225) = 3.6, p = 0.0003), EEAG, Words: *F*(9,225) = 23.6, p < 0.0001, EEAG, Scenes (*F*(9,225) = 13.9, p < 0.0001)] and the right [EEAG, LinearB (*F*(9,225) = 3.4, p = 0.0006, EEAG, Words *F*(9,225) = 18.0, p < 0.0001, EEAG, Scenes (*F*(9,225) = 11.4, p < 0.0001)] hemispheres. Thus, the activation profile under the linear B script, words and scenes conditions compared to their scrambled version differed from that under the EEAG condition, although some regions exhibited the same level of activation (Figure 3A). The plot of activations under the EEAG condition against the activations under each of the other conditions suggests that a positive linear relationship exists only between the objects and EEAG conditions (Figure 3B).

To verify that the relationship observed at the group level between the activation profiles elicited by the objects and EEAG was not determined by outliers, the correlation of the activation profiles in 10 regions in response to the EEAG and objects (compared to their scrambled version) was computed in each hemisphere separately for each subject. Then, the resulting Pearson’s correlation coefficients of each subject were Fisher z-transformed and analysed using a univariate two-sided t-test that was significant in both the left (t(25) = 7.6, p < 0.0001) and right (t(25) = 4.8, p < 0.0001) hemispheres. This finding confirms that both EEAG and object perception recruit the ventral pathway in a similar way in both hemispheres.

The proximity of the ventral visual cortex responses to these two visual categories was likely elicited by the global visual organization of EEAG, which triggered processes comparable to those required for object recognition. Among the regions activated by the perception of both EEAG and objects, O3-1 and O3-2 corresponded to the so-called LO functional region^20^. LO is defined as the brain area showing the greatest activation while viewing a known or novel object compared to its scrambled version^20,21^. The involvement of this region further supports the hypothesis that the processes involved in object recognition are involved in the perception of EEAG. This hypothesis is consistent with the fact that LO has been proven to be sensitive to the shape, but not the semantics, of objects^22,23^. Thus, although the similarity between the profiles of the objects and EEAG does not demonstrate that EEAG were previously used as symbols, modern brains perceive these EEAG as coherent visual entities to which symbolic meaning can be attached. Given that it has recently been argued that the anatomy of cortical areas belonging to the visual ventral cortex have been moderately impacted by the evolution of the brain from the earliest hominins to *Homo sapiens*^24,25^, it seems reasonable to speculate that the present results also apply to early modern humans and archaic hominins. Considering that EEAG were produced deliberately and had no apparent utilitarian function, our results are consistent with the hypothesis that these graphic productions had a representational purpose and were used and perceived as icons or symbols by modern and archaic hominins.

At the regional level, only one area, i.e., the occipito-temporal region of the left hemisphere or FUS4, was activated by both EEAG and tasks involving symbolic or iconic contents (objects, linear B, and words). This region corresponds to the so-called visual word form area (VWFA)^26,15^. This region was activated by the perception of both objects, i.e., the linear B script and words, which is consistent with the proposal that the role of the left FUS4 is not restricted to word recognition^27–29^. As proposed by Vogel, FUS4 processing extends to the processing of complex visual perception with a statistical regularity and a “groupable” characteristic (i.e., perceived as a whole)^28^. Crucially, our results suggest that these features are found not only in words, images of objects and strings of symbols but also in EEAG, supporting their potential representational value.

Remarkably, FUS4 was also the only region where the leftward asymmetry survived the FDR correction under the EEAG condition (mean = 0.12, t(25) = 3.17, p = 0.04 FDR corrected, one-sample two-sided t-test). As previously described^16^, word presentation causes the largest leftward asymmetry likely due to hemisphere high-order language areas top-down processing leading to right FUS4 deactivation (Table 1). A significant, but lower intensity, leftward asymmetry was also present during the perception of objects and the linear B script (Figure 4A, all p’s < 0.05 corrected, one-sample two-sided t-tests). The leftward asymmetries measured during the EEAG, linear B script and object presentations did not significantly differ (all p’s > 0.10, post hoc t-tests) but were significantly larger than those measured during the scene perception (all p’s < 0.01) and significantly smaller than those triggered in the words minus scrambled words contrast (all p’s < 0.0001). To ensure that this result was not determined by variation under the scrambled pictures conditions, we confirmed that the perception of the scrambled pictures of each category did not result in any significant asymmetry in this region (compared to the fixation dot, see methods, all p’s > 0.10 uncorrected, Figure 4B).

**Figure 4.**
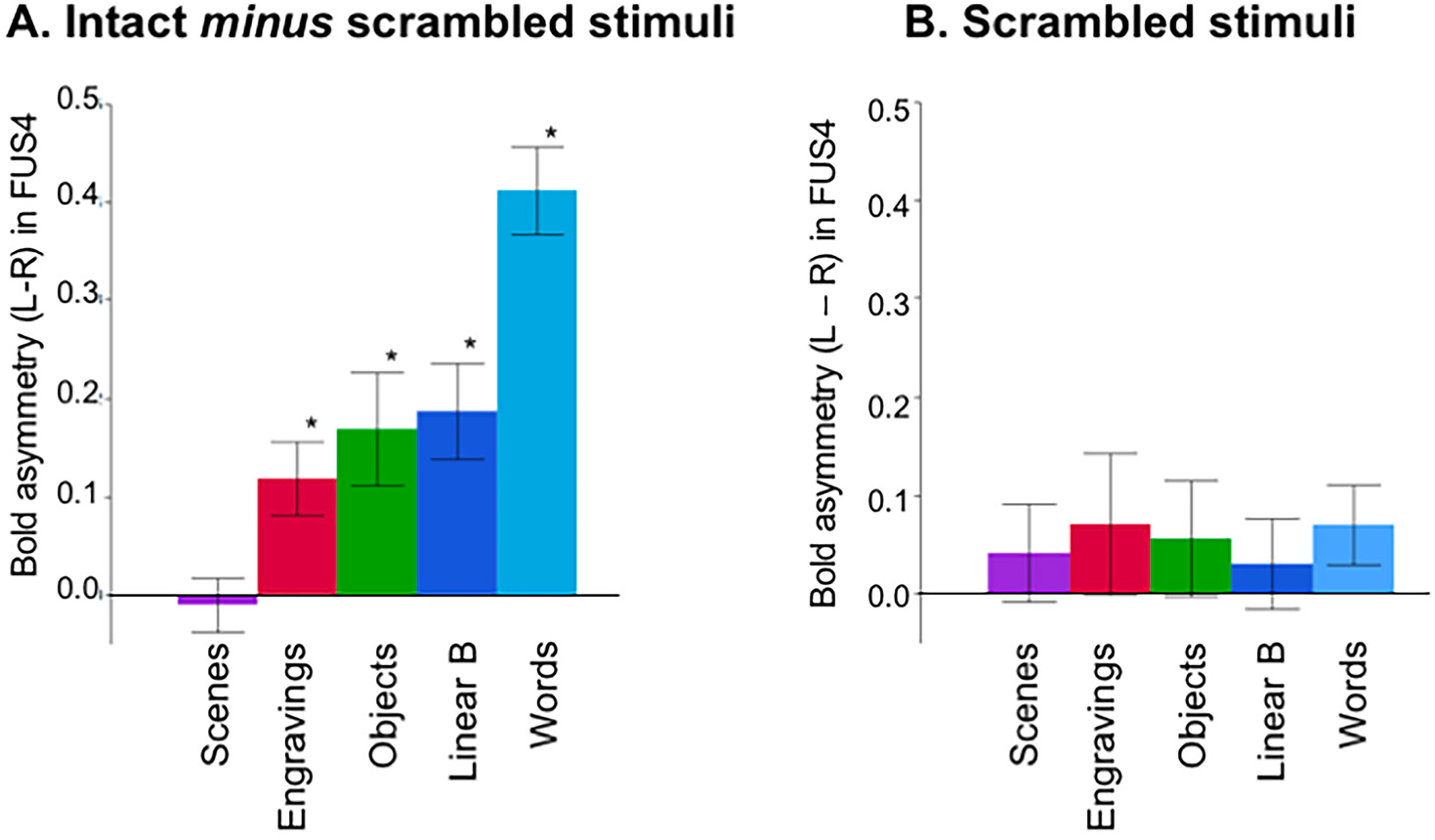
Asymmetries of BOLD signal (left minus right) in FUS4 under the five conditions **A.** Asymmetries measured in the contrast of intact minus scrambled stimuli. **B.** Asymmetries measured during the perception of the scrambled version of each category of stimuli (*: p < 0.05, one-sample t-test, FDR corrected. Error bars represent the standard error of the mean).

Importantly, the left hemisphere hosts language in most right-handed individuals. More generally, the left hemisphere is dedicated to cultural artifacts processing in most humans^30^. Under this framework, the leftward asymmetry observed in FUS4 during the perception of EEAG suggests that EEAG share some representational features with both objects and the linear B script that are not present in scenes, which did not show any asymmetry (p = 0.72, one-sample two-sided t-test). The trend of gradually decreasing leftward asymmetries in FUS4 from words, objects, linear B characters, EEAG to scenes may reflect the meaning conveyed by the different categories of stimuli; i.e., the more a stimulus is identified as conveying meaning, the more important the leftward asymmetry (Figure 4). This finding can be considered additional evidence supporting that EEAG are recognized as forms that could have been used by a human culture in the past to store and transmit coded information. Thus, the human brain perceives these graphic entities as having significant regularities to which semantic information can be connected. The present results support the as sumption that this type of production was not exclusively associated with anatomically modern humans.

## Materials and Methods

### Participants

The study was approved by the “Sud-ouest outremer III” local Ethics Committee (N° 2016/63). Twentysix healthy right-handed adults (mean age ± SD: 21.7 ± 2.5 years, 15 women) with no neurological history were included after providing written informed consent to participate in the study. The participants had a mean educational level of 15 years of schooling after first grade. One participant was excluded from the analysis due to technical issues in the fMRI images.

### Data acquisition

#### Stimuli

Our experimental protocol included five categories of stimuli, and each category was compared to its scrambled version (Figure 1).

The stimuli of interest, labelled EEAG, consisted of 50 tracings of abstract patterns created with stone tools on a variety of media that were discovered at archaeological sites dating from the Lower, Middle and Early Upper Palaeolithic eras (see Table S1 in supplementary material). We used tracings in which the engraved lines were rendered in white on a grey background and the outline of the object was rendered in light grey. The thickness of the lines on the tracings reflects the width of the engraved lines on the original piece. These stimuli were 700 * 480 pixels in size.

The pictures of objects (256 * 256 pixels) were represented by/consisted of nameable human-made artefacts depicted in different shades of grey. The stimuli were used to determine whether the neural bases of the engravings’ perception can be distinguished according to their levels of perceptual organization and the meaning they convey. The lowest level of organization was represented by the scrambled pictures, which were designed to serve as a control for the processing of each condition. These stimuli were constructed by randomly scrambling the other pictures into 32 to 124 squares depending of the initial size of the original picture using MATLAB software (MathWorks, Inc., Sherborn, MA, USA). This treatment removes any perceptible organization from the images^19^. These stimuli were included as matched reference stimuli and used to subtract non-specific activations linked to the processing of any type of visual information (stimuli were equal in global luminance and size).

The corpus of linear B consisted of 50 images of six characters strings written in white on a grey background. These images were 653 * 114 pixels in size.

The word stimuli were bi-syllabic and composed of six letters in lowercase on a grey background. The word stimuli were nouns referring mostly to a concrete notion (80%). The pictures of words had a size of 256 * 64 pixels. The word frequency was 15.5 ± 3 (SD) according to the lexical data Lexicon 3^30^. Low frequency words were chosen since these words activate ventral occipito-temporal regions more strongly than frequent words^31^.

The outdoor scenes were 256 * 256 pixels pictures of scenes, including the landscapes of mountains, beaches, or cities.

#### fMRI acquisition

The imaging was performed using a Siemens Prisma 3 Tesla MRI scanner. The structural images were acquired with a high-resolution 3D T1 weighted sequence (TR = 2000 ms, TE = 2.03 ms; flip angle = 8°; 192 slices; and 1 mm^3^ isotropic voxel size). In addition B0 maps were acquired to correct for geometric distortion (two acquisitions of 18s repeated four times). The functional images were acquired with a whole-brain T2*-weighted echo planar image acquisition (T2*-EPI Multiband x6, sequence parameters: TR = 850 ms; TE = 35 ms; flip angle = 56°; 66 axial slices; and 2.4 mm^3^ isotropic voxel size). The functional images were acquired in three sessions. The experiment presentation was programmed in E-prime software (Psychology Software Tools, Pittsburgh, PA, USA). The stimuli were displayed on a 27” screen. The participants viewed the stimuli through the rear of the magnet bore via a mirror mounted on the head coil.

#### Experimental protocol

The acquisition was organized in 3 sessions. The participants performed a 1-back task involving pictures of objects, scenes, and words and their scrambled versions in one session.

During the first run, the stimuli consisted of 234 distinct pictures presented in greyscale in a bitmap format belonging to one of the following three categories: scenes, objects, and words in their intact and scrambled versions. There were 39 different pictures per category. The block-designed paradigm consisted of 18 blocks lasting 12.75 s (6 blocks per category and their scramble version). Each of the 15 stimuli within a block was displayed for 300 ms, including two repetitions, and the participants were asked to detect the stimuli. A fixation point was displayed for 12.75 s every two blocks.

During the second and third runs, the subjects performed the same task, and the stimuli were the engravings, linear B strings and their scrambled version. Each run lasted for 4 min and 12 s with a random order of presentation and comprised 12 experimental blocks of 13.6 s interleaved with 7 fixation blocks of 12.75s. Each experimental block contained 11 stimuli and 2 repetitions with a 200 ms presentation time for each stimulus and an interstimulus interval of 1037 ms.

#### Image processing

Image analysis was performed using SPM12 software. The T1-weighted scans of the participants were normalized to a site-specific template (mean of 500 age-matched normal volunteers) matching the MNI space using the SPM12 “segment” procedure with the default parameters. To correct for subject motion during the fMRI runs, the 192 EPI-BOLD scans were realigned within each run using a rigid-body registration. Then, the EPI-BOLD scans were rigidly registered structurally to the T1-weighted scan. The combination of all registration matrices allowed for the warping of the EPI-BOLD functional scans to the standard space with a trilinear interpolation. Once in the standard space, a 6-mm FWHM Gaussian filter was applied.

#### Regional analysis

Within the selected hROIs, the activation profiles across the regions were calculated for each condition contrast (engravings minus scramble, objects minus scramble, scenes minus scramble, linear B minus scramble and words minus scramble) for each of the 26 participants.

The objects, linear B and scenes were compared to the engravings using repeated- measures 3-way ANOVA in each hemisphere (i.e., 2 (conditions) x 10 (hROIs)). In addition, Pearson’s correlation coefficients between the profiles of the engravings and objects conditions were computed for each subject and Fisher z-transformed. Then, the 26 Z-scores were analysed using a univariate two-sided t-test to determine whether the profile of objects was correlated to the profile of engravings. This analysis was conducted in each hemisphere separately.

A list of the regions activated by the objects, words, linear B, and scenes compared to their respective scrambled counterparts is presented in Table S3 to S6 as supplementary material.

Since a leftward asymmetry in the ventral route is typical of the processing of semantic and verbal material^16,27^, we calculated the left minus right BOLD values in the selected hROIs during the EEAG presentations to identify the hROIs with significant leftward asymmetries (one-sample t-test, FDR corrected). Then, we compared the identified hROI(s) asymmetries across all conditions with one-way ANOVA.

## Acknowledgements

This research was supported by a grant from IdEx Bordeaux / CNRS (PEPS 2015). The authors thank GinesisLab (Programme Labcom 2016, ANR 16LCV2-0006-01) for their help in data management and data processing and Carole Peyrin for providing some of the stimuli used.

## Author contributions

EM, NTM and FE conceived the study. EM, MS, GJ, and AM designed the study. EM, MS, and SC performed the experiment, EM, MS, SC, NTM, MJ, and BM carried out the analyses, EM, NTM and FE wrote the manuscript with contributions from AM, BM, and GJ.

## Supplementary Materials

**Table S1.** Contextual and descriptive data on early engravings used as visual stimuli. CC: Criss-cross; CH: Cross-hatching; CHB: Cross hatched band; Ch/Qt: Charentian – Quina type; CL: Curved lines; CO: Concentric lines; EUP: Early Upper Paleolithic; FL: Fan-like; HB: Hatched band; LP: Lower Paleolithic; MP: Middle Paleolithic; M: Mousterian; MSA: Middle Stone Age; OES: Ostrich egg shell; PAR: Parallel lines; PM: para-Micoquian; PMK: Parallel marks; SPL: Sub-parallel lines; ZZ: Zigzag.

| Site | Region | Material | No. of objects | Cultural Attribution | Age (kyr BP) | Description | Reference |
| --- | --- | --- | --- | --- | --- | --- | --- |
| **Africa** |  |  |  |  |  |  |  |
| Blombos | South Africa | Ochre; Bone | 18 | MSA | 100-75 | CH; PAR; CC | d'Errico et al. 2001; Henshilwood et al. 2002, 2009 |
| Diepkloof | South Africa | OES | 7 | MSA | 65-55 | SPL; CL; CHB | Texier et al. 2010 |
| Klasies River | South Africa | Ochre | 1 | MSA II | 100-85 | SPL | d'Errico et al. 2012 |
| Klein Kliphuis | South Africa | Ochre | 1 | MSA | 80-50 | SPL; CH | Mackay and Welz 2008 |
| Klipdrift Shelter | South Africa | OES | 1 | MSA | 65–59 | CH; CC; SPL | Henshilwood et al. 2014 |
| Pinnacle Point | South Africa | Ochre | 1 | MSA | 100 | SPL | Watts 2010 |
| Palmenhorst | Namibia | Stone | 1 | MSA | Undated | CH; CC; PAR | Henshilwood and d'Errico 2011 |
| Sibudu Cave | South Africa | Ochre | 3 | MSA | 77-58 | PAR; FL | Hodgskiss 2014 |
| **Europe** |  |  |  |  |  |  |  |
| Bacho Kiro | Bulgaria | Bone | 1 | MP-M | >47 | ZZ | Marshack 1976, 1982 |
| Bilzingsleben | Germany | Bone | 2 | LP | 412-320 | SPL; FL | Mania and Mania 1988 |
| Ermitage | France | Bone | 1 | MP-M | >31 | PAR; SPL | Pradel and Pradel 1954; d'Errico 1998 |
| Gorham's Cave | Gibraltar | Bedrock | 1 | MP-M | >39 | CH | Rodríguez-Vidal et al. 2014 |
| Klik-Koba | Crimea | Flint | 1 | MP-PM | 35-37 | SPL | Stepanchuk 2006, Majkić et al. 2018 |
| Kozarnika | Bulgaria | Bone | 1 | LP | 900 | PMK | Sirakov et al. 2010; Ivanova 2009; Guadelli and Guadelli 2004 |
| Krapina | Croatia | Bone | 1 | MP | 130 | PMK | Frayer et al. 2006 |
| La Ferrassie | France | Bone | 1 | MP-M | 65 | PMK | Capitan and Peyrony 1912; Zilhao 2007 |
| Mamontovaya Kurya | Russia | Tusk | 1 | MP-M | 36 | SPL | Pavlov et al. 2001; Svendsen, Pavlov 2003 |
| Pešturina | Serbia | Bone | 1 | MP-Ch/Qt | 95–64 | SPL | Majkić et al. 2017 |
| Temnata Dupka | Bulgaria | Stone slab | 1 | MP | 50 | PAR | Cremades et al. 1995; Tsanova 2006 |
| **Middle East** |  |  |  |  |  |  |  |
| Qafzeh | Israel | Flint | 1 | MP-M | 100 | PAR | Hovers et al. 1997 |
| Quneitra | Israel | Flint | 1 | MP-M | 50 | CO | Goren-Inbar 1990; Marshack 1996; d'Errico et al. 2003 |
| **Asia** |  |  |  |  |  |  |  |
| Shuidonggou | China | Stone | 1 | EUP | 30 | PAR | Peng et al. 2012 |
| Trinil | Indonesia | Shell | 1 | LP | 540-430 | ZZ; PAR | Joordens et al. 2015 |

**Table S2.** Abbreviations and MNI Coordinates of the mass center of the ten hROIs of the Aicha atlas activated by the perception of engravings minus scramble contrast.

| hROI | Abbreviation | Left hemisphere | Right hemisphere |
| --- | --- | --- | --- |
|  |  | MNI coordinates | MNI coordinates |
| G_Fusiform-2 | FUS2 | -35 -26 -23 | 38 -25 -24 |
| G_Fusiform-3 | FUS3 | -37 -32 -24 | 37 -31 -24 |
| G_Fusiform-4 | FUS4 | -43 -50 -17 | 44 -46 -18 |
| G_Temporal_Inf-4 | T3-4 | -50 -61 -8 | 54 -58 -11 |
| G_Fusiform-5 | FUS5 | -31 -50 -12 | 32 -47 -11 |
| G_Temporal_Inf-5 | T3-5 | -45 -64 6 | 49 -58 4 |
| G_Occipital_Inf-1 | O3-1 | -48 -69 -4 | 50 -60 -9 |
| G_Occipital_Inf-2 | O3-2 | -45 -71 -7 | 47 -65 -7 |
| G_Occipital_Mid-2 | O2-2 | -37 -77 14 | 41 -73 12 |
| G_Occipital_Lat-3 | OLat3 | -40 -84 -12 | 43 -81 -12 |

**Table S3.** Mean BOLD value of the hROIs activated in the Words minus scrambled words contrast (p< 0.05 uncorrected)

| Activation of words minus scrambled words |  |  |  |  |  |  |  |  |
| --- | --- | --- | --- | --- | --- | --- | --- | --- |
|  | Left hemisphere |  |  |  | Right hemisphere |  |  |  |
|  | Mean BOLD | SD | t | p | Mean BOLD | SD | t | p |
| G_Fusiform-1 | 0.11 | 0.02 | 4.40 | 0.0002 |  |  |  |  |
| G_Fusiform-2 | 0.07 | 0.03 | 2.27 | 0.0323 |  |  |  |  |
| G_Fusiform-4 | 0.25 | 0.06 | 4.22 | 0.0003 |  |  |  |  |
| G_Temporal_Sup-4 | 0.19 | 0.04 | 4.42 | 0.0002 |  |  |  |  |
| G_Temporal_Mid-4 | 0.14 | 0.04 | 3.21 | 0.0036 |  |  |  |  |
| G_Temporal_Inf-3 | 0.18 | 0.05 | 3.50 | 0.0018 |  |  |  |  |
| S_Sup_Temporal-4 | 0.32 | 0.05 | 6.13 | <.0001 |  |  |  |  |
| S_Sup_Temporal-3 |  |  |  |  | 0.19 | 0.05 | 3.48 | 0.0019 |

**Table S4.**
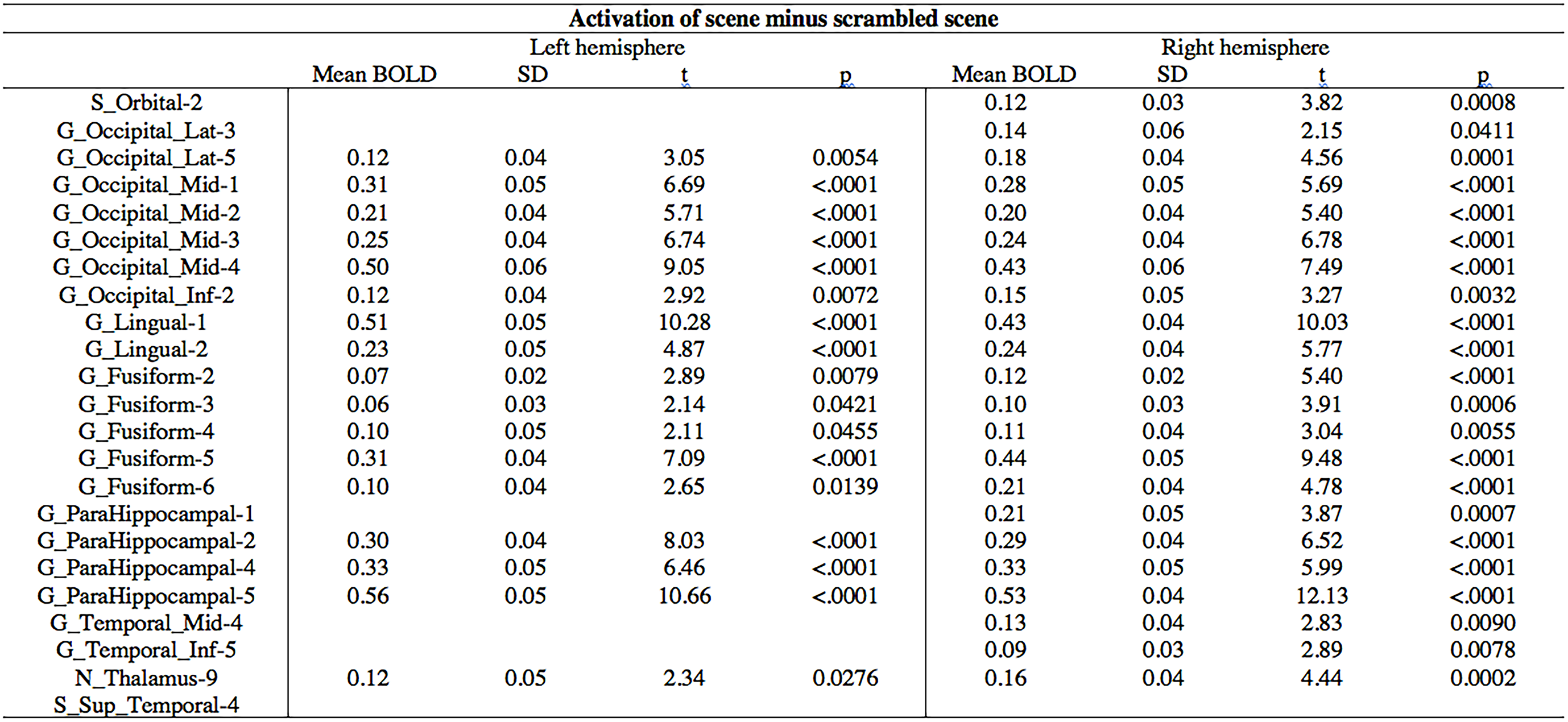
Mean BOLD value of the hROIs activated in the scenes minus scrambled scenes contrast (p< 0.05 uncorrected)

**Table S5.**
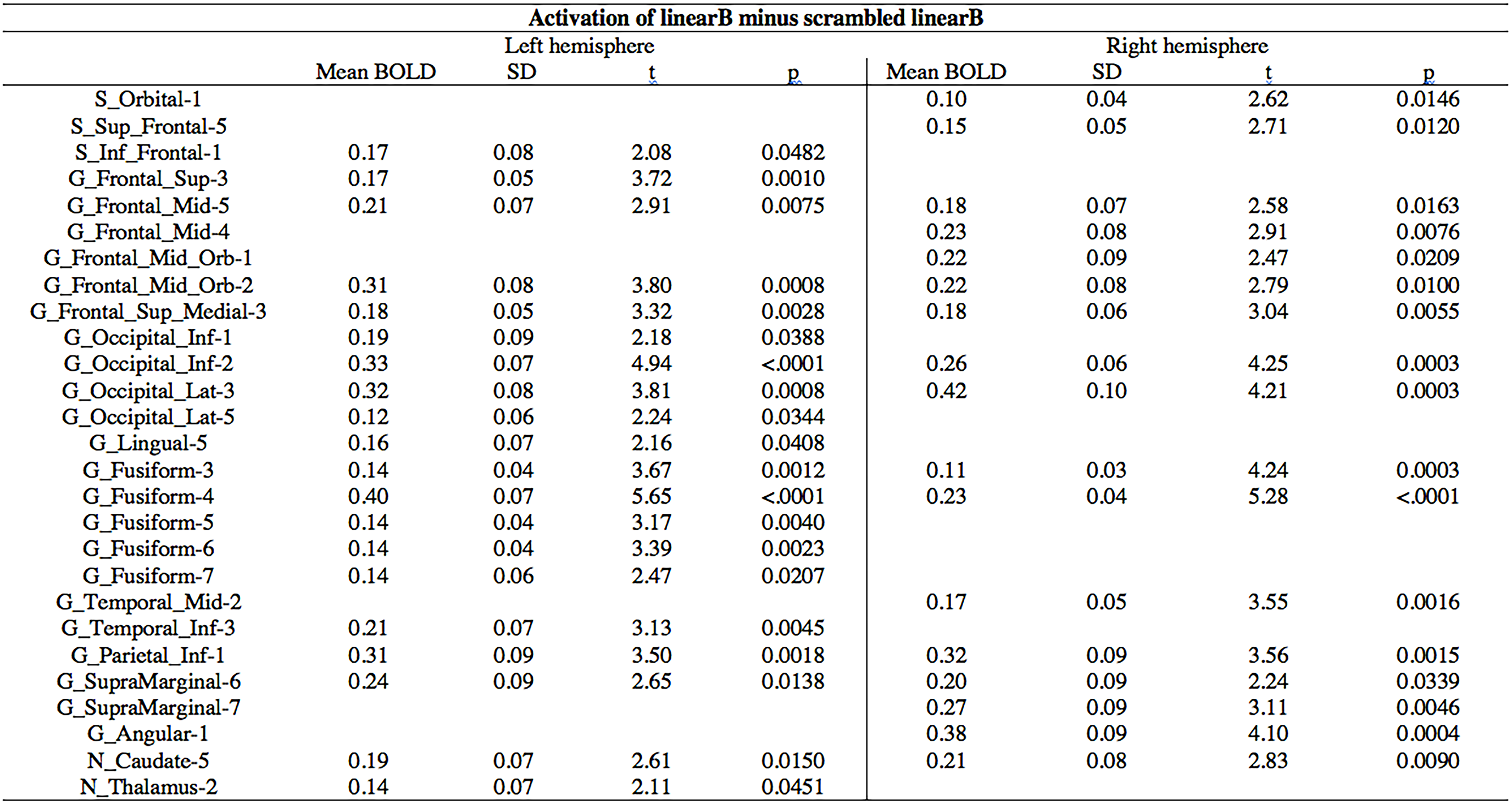
Mean BOLD value of the hROIs activated in the linear B minus scrambled linear B contrast (p< 0.05 uncorrected)

**Figure S1.**
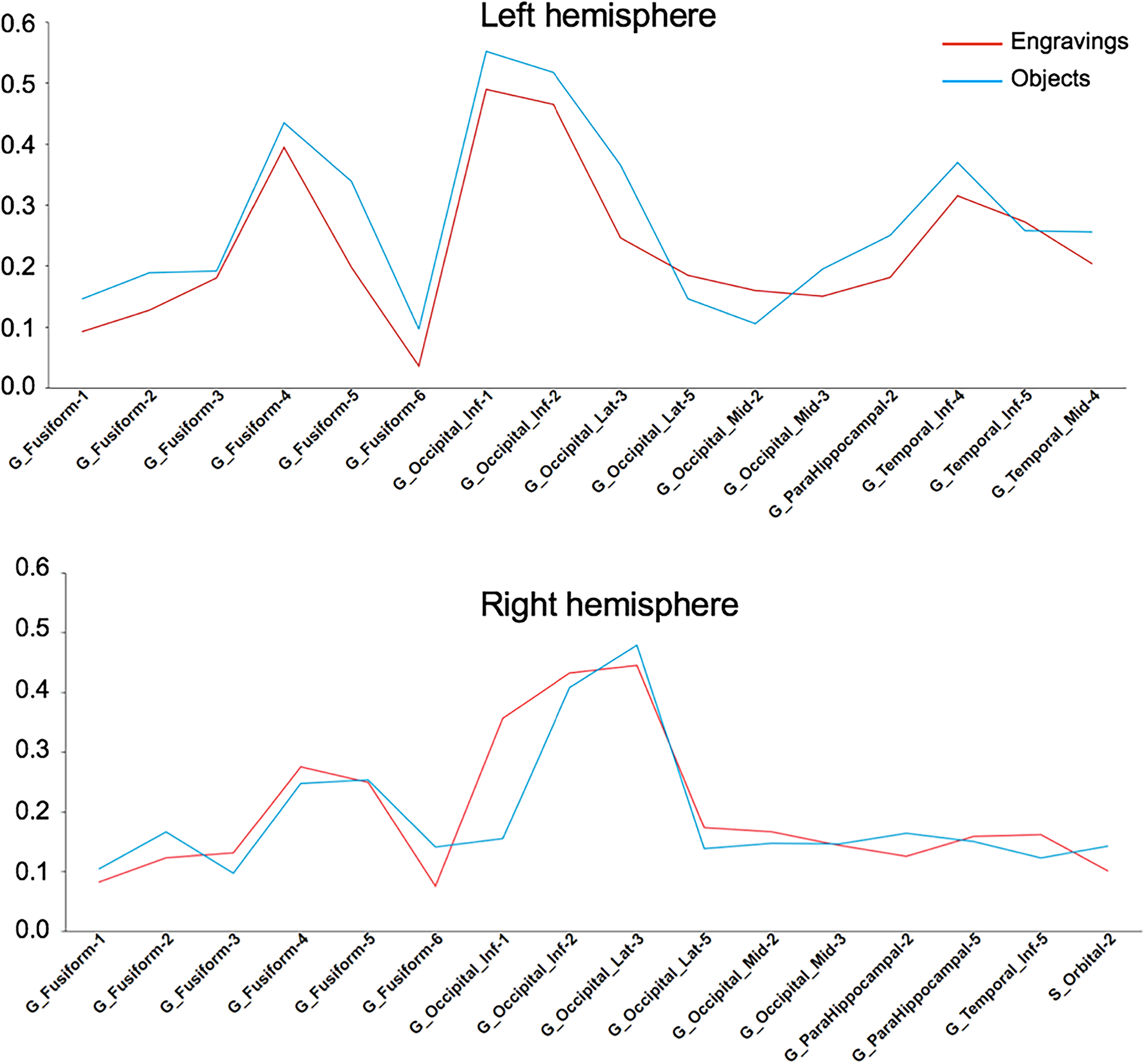
Descriptive analysis of the activation profiles under the Object and EEAG perception conditions of the 16 hROIs of the ventral route activated by object perception. No regions x conditions interaction was significant neither in the left nor in the right hemisphere (F(15,375) = 1.25, p = 0.23 and F(15,375) = 1.15, p = 0.31, respectively).

